# Single-Molecule Activity Profiling of Glycosidase Proteoforms Using Water-Soluble Fluorogenic Probes

**DOI:** 10.64898/2026.05.22.727124

**Authors:** Akihiro Kaneko, Kyohhei Fujita, Takako Uchida, Tadahaya Mizuno, Takumi Iwasaka, Hiroyuki Kusuhara, Yasuteru Urano, Toru Komatsu

## Abstract

In this study, we report a series of highly water-soluble fluorogenic probes for glycosidases useful for microdevice-based single-molecule enzyme activity profiling (SEAP). The assay was able to detect trace glycosidase activities in blood with high sensitivity and with proteoform resolution. This platform revealed the potential of α-mannosidase as sensitive blood-based liquid-biopsy biomarkers for liver injury, highlighting the potential of this approach for activity-based diagnostics.

---

Glycosidases are enzymes that hydrolyze the glycoside bonds of polysaccharides and the sugar chains of glycoproteins, and their functional dysregulation is a hallmark of various diseases, including cancer and lysosomal storage disorders^1–3^. Although numerous studies have attempted to understand the relationship between enzymatic activity alterations and phenotypic changes, conventional biochemical assays—typically analyzing 10^6^-10^9^ enzyme molecules in bulk—provide only ensemble-averaged data. Such approaches often mask the heterogeneous behaviors of important proteoforms that are closely linked to pathological states^4^. While single-molecule enzyme activity profiling (SEAP) using microdevices has emerged as a powerful tool to overcome these limitations^4–10^, its application has been restricted by the limited availability of suitable fluorogenic probes, particularly for glycosidases.

In this study, we report the development of novel glycosidase probes based on the sulfo-TokyoGreen (sTG) scaffold, which possesses high water solubility and superior fluorescence properties optimized for SEAP assays, facilitating the discovery of disease-specific proteoform landscapes (**Figure 1**)^5^. SEAP assays are performed by loading a diluted enzyme solution into a microfabricated chamber device consisting of 10^5^-10^6^ femtoliter-volume reactors. The activity of these single enzyme molecules is monitored using fluorogenic probes, allowing enzyme species with different catalytic activities—arising from distinct isozymes or proteoforms—to be distinguished based on differences in their reaction rates^4,11^. To achieve precise readout of enzyme activity, the fluorogenic probes must possess the superior fluorescence activating properties to report the weak signal from the turnover of single-molecule enzyme, and sufficient water solubility to prevent from the leakage of the fluorescent products into the surrounding oil layer of the microdevice^6^.

**Figure 1.**
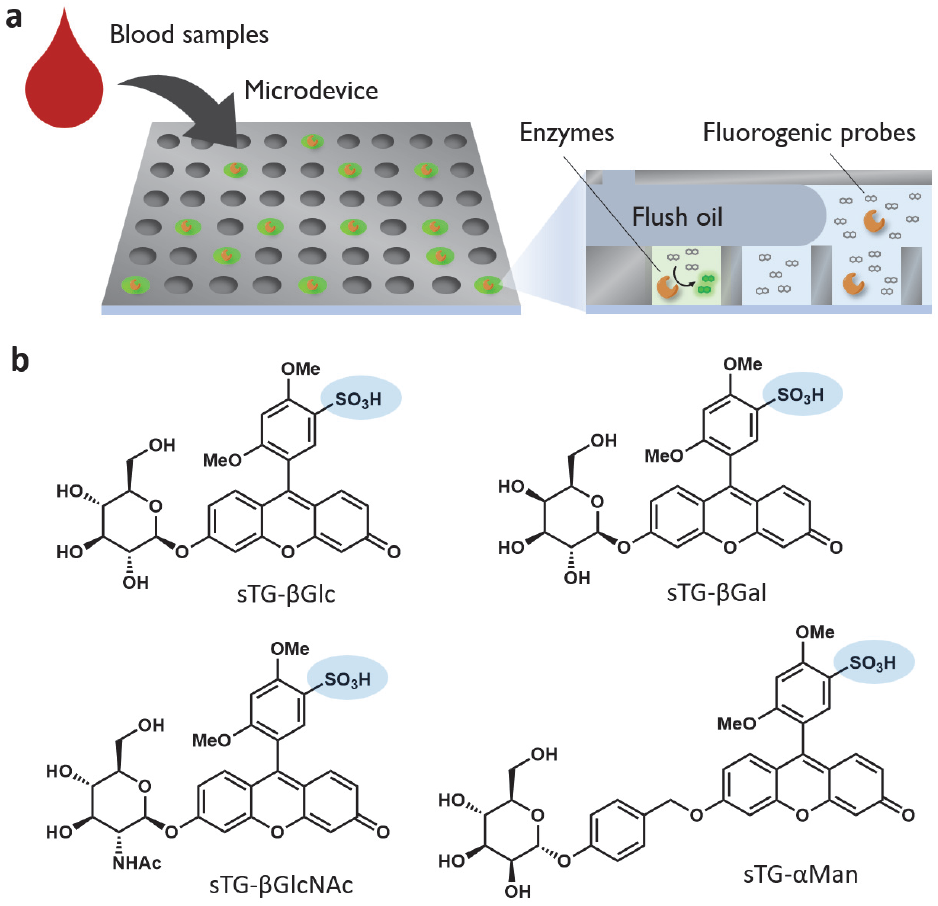
Design of fluorogenic probes for single-molecule glycosidase probes. (a) Scheme of single-molecule enzyme activity analysis of glycosidases. (b) Structures of fluorogenic probes synthesized based on sTG scaffold.

As discussed, none of the present probe scaffolds are suitable for detecting mammalian glycosidases in SEAP platforms, and we establish the design rationale of glycosidase probes based on sulfo-TokyoGreen (sTG) scaffold as a core fluorophore (**Figure 1b**). This scaffold, whose fluorescence activation is governed by a photoinduced electron transfer (PeT)^12^, has proven optimal for SEAP because the introduction of a sulfonic acid moiety suppresses leakage and ensures high retention within the femtoliter chambers^5^.

We designed and synthesized a series of sTG-based probes targeting various glycosidases, including β-glucosidase (sTG-βGlc), β-galactosidase (sTG-βGal), β-hexosaminidase (sTG-βGlcNAc), and α-mannosidase (sTG-αMan). For sTG-βGlc, sTG-βGal, and sTG-βGlcNAc, the corresponding monosaccharide was directly conjugated to the phenolic oxygen of sTG (**Scheme S1**). In contrast, for sTG-αMan, the directly modified glycoside bond was not stable enough, so we employed a *p*-hydroxybenzyl alcohol (PHBA; quinone methide-forming) linker between the sTG core and the α-D-mannoside moiety (**Scheme S2**). This self-immolative linker undergoes efficient 1,6-elimination to release the free fluorophore upon enzymatic cleavage^13,14^.To validate the performance of the synthesized sTG-based glycosidase probes, we first evaluated their fluorogenic responsiveness in bulk biochemical assays with recombinant enzymes. Upon incubation with their respective target glycosidases, all four probes exhibited significant increases in fluorescence intensity (**Figure 2**). For probes with direct carbohydrate substitution, sTG-βGlc, sTG-βGal and sTG-βGlcNAc, the fluorescence activation was more than 100-fold. For sTG-αMan, a probe with PHBA linker, the background fluorescence was slightly high, showing fluorescence activation of 27-fold. The relatively high background fluorescence might have attributed to the electron-donating PHBA contributed to the lower oxidation potential of excited state of protected xanthene, so the PeT process became less efficient^12^, but we considered that the fluorescence activation might be sufficient for the reliable quantification of enzymatic activities, especially when transitioning to the single-molecule level.

**Figure 2.**
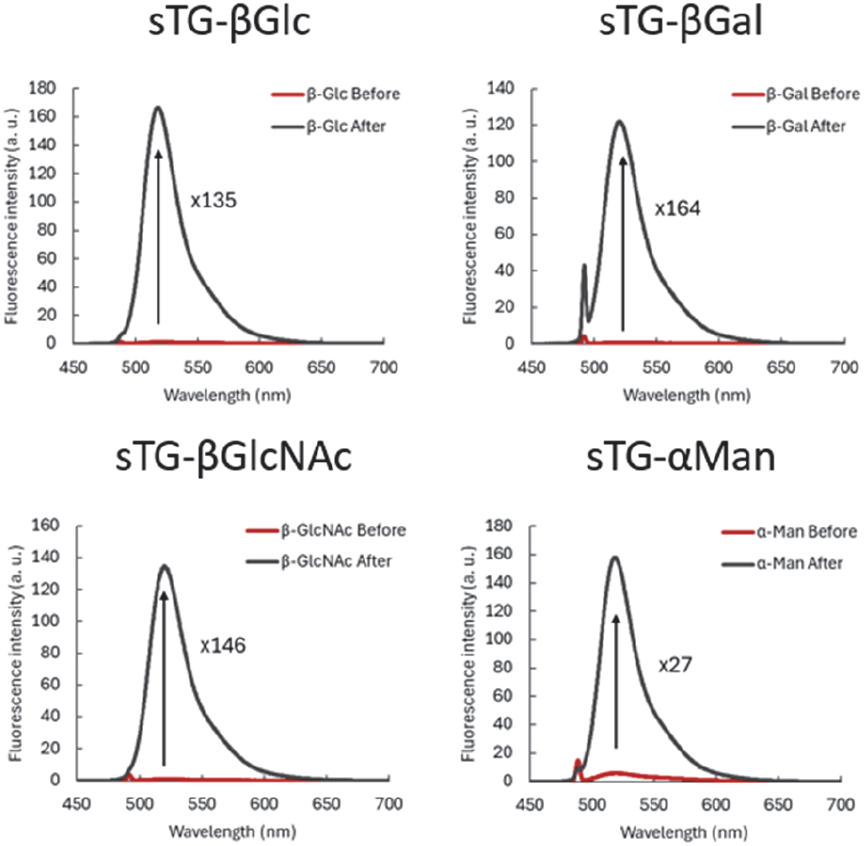
Fluorescence activation of glycosidase probes after reacting with enzymes. The probes (2 μM) were mixed with enzymes (0.2 mg/mL) in phosphate buffer (100 mM, pH 7.4) and incubated for 18 h. The fluorescence spectra before and after the enzymatic reaction were recorded under λ_ex._ = 490 nm.

A key requirement for SEAP is the orthogonal activation of multiple probes within a complex biological mixture. We next confirmed the substrate specificity of each probe by testing them against a panel of the four targeted glycosidases. Among them, GBA3, a cytosolic human β-glucosidase showed slightly promiscuous reactivity, reacting with sTG-βGal along with sTG-βGlc^15^, but each sTG-based probe was hydrolyzed primarily by its corresponding enzyme, showing the minimal cross-reactivity (**Figure 3**).

**Figure 3.**
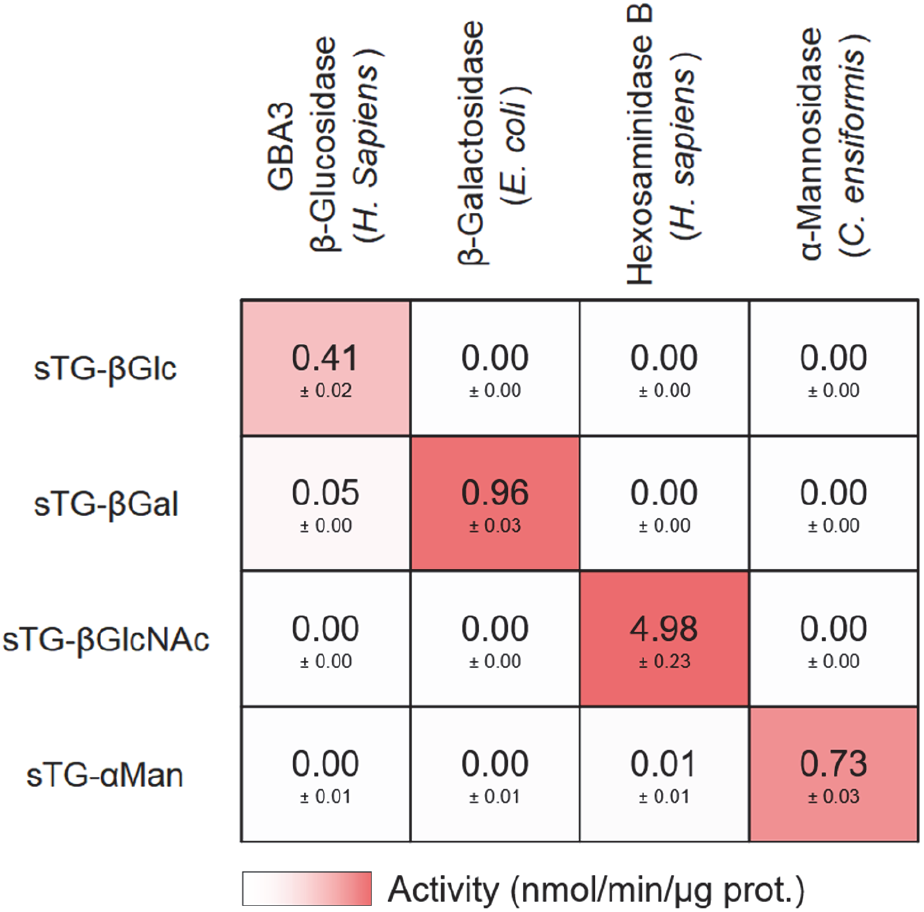
Orthogonality of glycosidase probes. Probes (10 μM) were reacted with enzymes (10 μg/mL) in phosphate buffer (pH 7.4) for 90 min. For the initial fluorescence increase rates, background fluorescence signal (from samples without enzymes) was subtracted, and the values were converted to the activity (nmol/min/μg prot.) using fluorescence signal of sTG as a reference. The values are mean ± S. D. (n = 4).

We then characterized the performance of the synthesized probes in microdevice-based single-molecule enzyme activity assay using β-galactosidase as model systems. lacZ (bacterial β-galactosidase) is commonly used as the reporter enzyme in the microdevice-based assays^16^. While the bacterial β-galactosidase has optimal pH around 7.4, most mammalian glycosidases function within the acidic environments of lysosomes, so the enzymatic assay should be performed in acidic conditions. To optimize the detection sensitivity, we adjusted the assay buffer to pH 6.5 to monitor the mammalian lysosomal glycosidases. This condition provides a strategic balance, maintaining sufficient enzymatic activity for lysosomal glycosidases while ensuring that a fraction of the sTG (p*K*_a_ = 6.4)^12^ fluorophore remains in its highly fluorescent phenolate form. In this condition, sTG-βGal was able to detect the single-molecule activity of mammalian β-galactosidase. In contrast, the present probes such as TG-βGal and resorufin-βGal cannot detect the activity since the partial protonation of the fluorophore increased the hydrophobicity, making the fluorescent product leaking out from the reaction chamber (**Figure S1**). Another fluorogenic probe, fluorescein di-β-galactopyranoside (FDG)^17^, was not leaky from the system due to the carboxylic acid of fluorescein, but the presence of two reaction sites made the activity report much slower than sTG-βGal. The result confirmed that the sTG serves as the ideal fluorophore to design the glycosidase probes to report the single-molecule activity of mammalian lysosomal glycosidases.

We then analyzed the single-molecule activities of glycosidases in mouse blood samples. Under the acidic condition (pH 6.5), we successfully detected single-molecule activities for all four targeted glycosidases in 1/1,000-diluted blood samples (**Figure 4a**). At neutral pH (7.4), the activity spots of α-mannosidase was detected with the comparable brightness with that in acidic pH, while other glycosidases exhibited negligible or weaker activity (**Figure 4a**). In the conventional multi-well plate-based bulk assay, a signal was observed only for β-hexosaminidase at 1/10-diluted plasma, but signals from other probes were not distinguishable from the background. This result confirmed the sensitivity of SEAP platform that enables the amplification of local fluorescence signals to detectable levels from a single-molecule turnover. Besides the sensitivity, a unique strength of SEAP is its ability to resolve the heterogeneity of functional proteoforms within a single sample. Analysis of the activity intensity distribution for β-hexosaminidase (by sTG-βGlcNAc) revealed a distinct multi-modal distribution, suggesting the simultaneous presence of at least four functional species with different catalytic turnover rates (**Figure 4c**).

**Figure 4.**
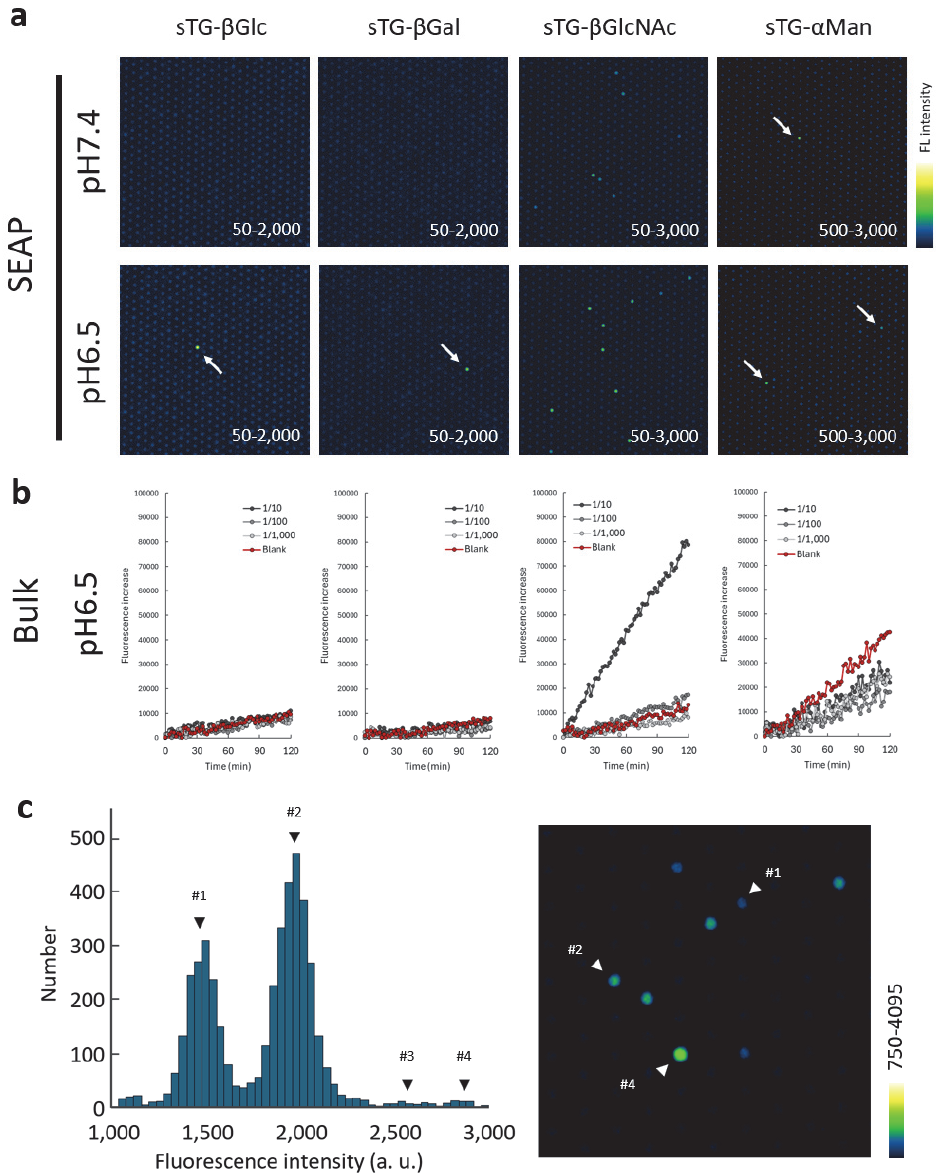
Detection of glycosidase activities in mouse blood samples using SEAP platform. (a) Fluorescence images of microdevice loaded with probe (30 μM) and blood samples (1/1,000) in sodium phosphate buffer (100 mM, pH 6.5) containing Triton X-100 (250 μM) or HEPES-Na buffer (100 mM, pH 7.4) containing CaCl_2_ (1 mM), MgCl_2_ (1 mM), DTT (100 μM) and Triton X-100 (250 μM). Images were acquired after incubating at 25°C for 18 h. (b) Fluorescence increase of fluorescent probes (10 μM) and blood samples (1/10-1/1,000) in bulk 384-well plate-based assay. The assay was performed in sodium phosphate buffer (pH 6.5) containing 0.1% CHAPS. In blank condition (red line), blood sample was not included. (c) Histogram showing the fluorescence distribution for sTG-βGlcNAc in (a). The spot with the corresponding activity are shown in magnified image.

Next, to evaluate the potential of our platform for activity-based diagnostics, we performed blood-based biomarker discovery study using two distinct mouse models of liver injury: the thioacetamide (TAA) model (inducing direct hepatocyte toxicity) and the 4,4’-methylene dianiline (MDA) model (inducing biliary-targeted damage)^6,18^. We analyzed the alterations in single-molecule glycosidase activity profiles associated with these pathological states (Figure 5). Our SEAP platform identified significant shifts in the enzymatic landscape of liver injury models (**Figure 5**). Specifically, we observed a dramatic increase in the number of active α-mannosidase molecules in the blood of TAA-treated mice compared to healthy controls. Individual sample evaluations (n = 4 for healthy controls and n = 6 for TAA-treated mice) confirmed this marked elevation, showing the potential of α-mannosidase as a potential biomarker for hepatocyte damage **(Figure 5b**). Furthermore, we detected a specific shift in the proteoform distribution of β-hexosaminidase. In both TAA and MDA models, the population of high-activity clusters (identified as Spot #2, #3, and #4) significantly increased (**Figure 5c, 5d**).

**Figure 5.**
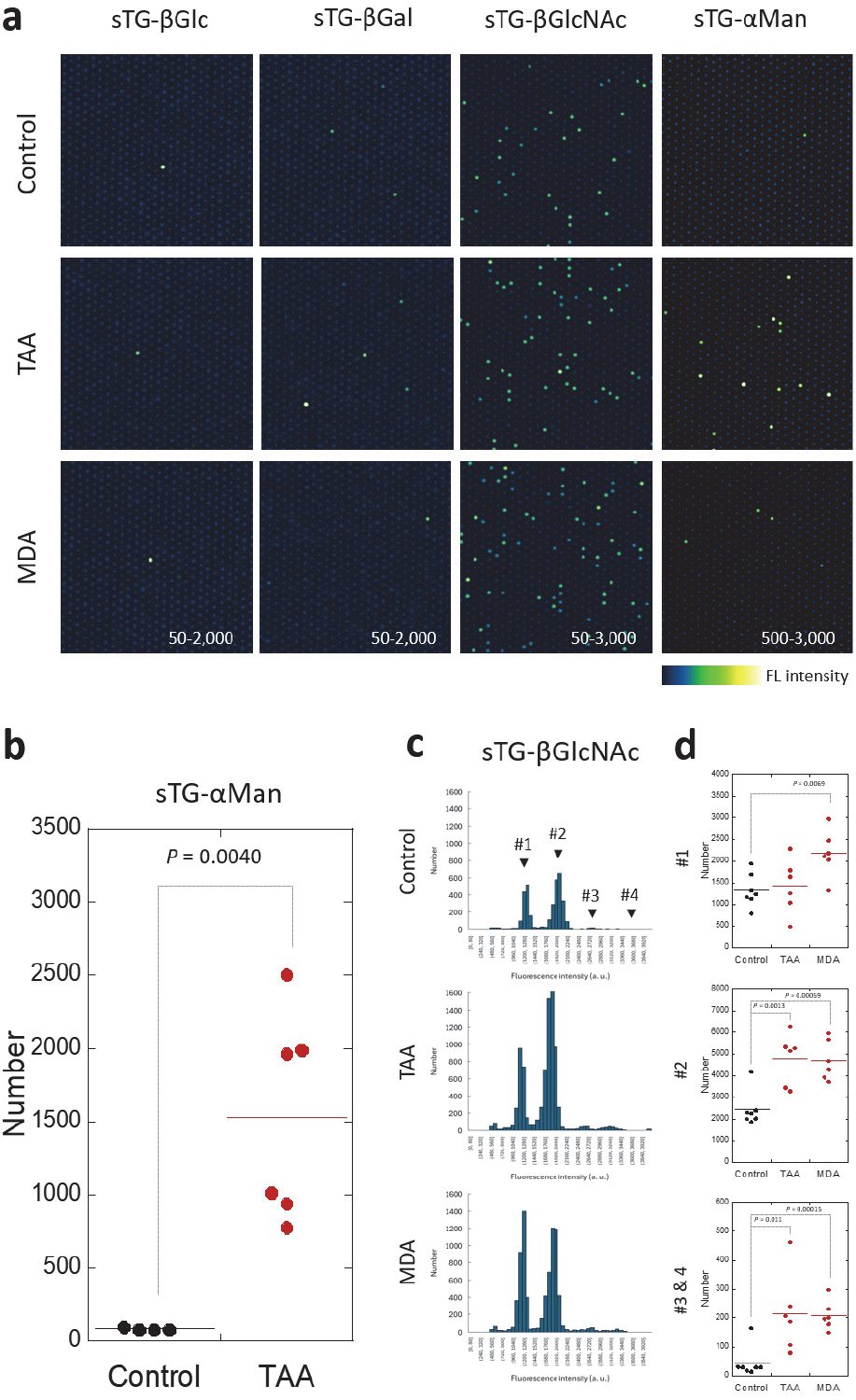
Blood-based biomarkers of liver damage. (a) Fluorescence images of microdevice loaded with probe (30 μM) and blood samples (1/300) in sodium phosphate buffer (pH 7.4 or 6.5) containing Triton X-100 (250 μM) and incubated for 18 h. (b) Quantification of the number of spot in the analysis of sTG-αMan in healthy control and TAA-treated mice. *P* value was calculated for Student’s t-test (n = 4 for healthy control, n = 6 for TAA). (c) Histograms showing the fluorescence distribution for sTG-βGlcNAc in (a). (d) Quantification of the number of spot of spot 1-4 in (c). *P* value was calculated for Student’s t-test (n = 7 for healthy control, n = 6 for TAA and MDA).

For spot #1, the increase was observed only for MDA-treated model. While both TAA- and MDA-treatment results in the damage to the liver, their mechanism of action are different; TAA triggers the hepatocellular damage, while MDA causes cholestasis-like behavior^6,18^. How the pattern differences can be connected to the pathological differences is still under investigation, but the results show that the detailed profiling of pathological proteoforms-which is lost in conventional bulk assays-provides a more nuanced understanding of disease progression and underscores the power of SEAP assays for the discovery of novel biomarker candidates.

In conclusion, we established a single-molecule enzyme activity profiling (SEAP) platform using water-soluble sTG-glycosidase probes to resolve pathological proteoforms in circulation that remain undetectable in conventional bulk assays. This technology facilitates the discovery of novel activity-based diagnostics, as demonstrated by identifying α-mannosidase and specific proteoform of β-hexosaminidase as a sensitive liquid biopsy biomarker for liver injury.

## Supporting Information

Methods, supplementary figures, and synthesis and characterization of probes.

## Acknowledgements

This work was financially supported by MEXT (20H04694, 21A303, 22H02217, 23K23484, 25K01911 and 25K2252), JST (FOREST [24012649]), and AMED (FORCE [22581634] and P-PROMOTE [25131640]). T. K. received support from the Naito Foundation, The Mochida Memorial Foundation for Medical and Pharmaceutical Research, the Chugai Foundation for Innovative Drug Discovery Science, the MSD Life Science Foundation, the Hoansha Foundation, and the University of Tokyo Gap Fund Program.

## Notes

### Competing Interest Statement

T. Mizuno and T. Komatsu are shareholders and advisors of Cosomil, inc.

