## Supplementary_Information for "Single-Molecule Activity Profiling of Glycosidase Proteoforms Using Water-Soluble Fluorogenic Probes"

**Contents**

**Methods**

**Supplementary figures**

**Supplementary methods for synthesis and characterization of compounds**

**Supplementary references**

### Methods

#### Materials

| REAGENT OR RESOURCE | SOURCE | IDENTIFIER |
|---|---|---|
| β-glucosidase from almonds | Sigma-Aldrich | Cat. #G4511<br>Lot #0000365877 |
| GBA3 from *Homo sapiens* | R&D Biosystems | Cat. #5969-GH<br>Lot #TLY042561 |
| β-galactosidase from *Escherichia coli* | Sigma-Aldrich | Cat. # G6008<br>Lot #0000505921 |
| GLB1 from *Homo sapiens* | R&D Biosystems | Cat. #6464-GH<br>Lot #DADI0322111 |
| Hexosaminidase B from Homo Sapiens | R&D Biosystems | Cat. #8907-GH<br>Lot #DEZE0225051 |
| α-Mannosidase from *Canavalia ensiformis* | Sigma-Aldrich | Cat. #M7257<br>Lot #0000354474 |

#### Instruments

NMR spectra were recorded on a JEOL JNM-LA400 instrument at 400 MHz for $^{1}$H NMR and at 100 MHz for $^{13}$C NMR. Mass spectra (MS) were measured with a JEOL JMS-T100LC AccuToF (ESI). LC-MS analyses were performed on a Waters Acquity UPLC (H Class)/QDa quadrupole MS analyzer or Acquity UPLC (H Class)/Xevo TQD quadrupole MS/MS analyzer equipped with an Acquity UPLC BEH C18 column (Waters). Column chromatography using silica gel was performed on an MPLC system (Yamazen Smart Flash EPCLC AI-5805 (Tokyo, Japan)). Reversed-phase MPLC purification was performed on an Isolera One (Biotage) equipped with a SNAP Ultra C18 30 g (Biotage).

**Digital enzyme assay in microdevice**

Digital enzyme assays were performed using commercially available microdevices (Simoa disk; Quanterix). A 35 μL of mixture of enzyme and reagents in buffer was loaded into the microdevice by manual pipetting. Then, 70 μL of FC-70 (Sigma-Aldrich) was introduced into the device to flush out excess reaction mixture. The enzymatic activity in the chambers was measured using an epifluorescence microscope (Ti2, Nikon) equipped with a 20× dry objective lens (Plan Apo 20×), an sCMOS camera (ORCA-Fusion C14440, Hamamatsu Photonics), a white LED illumination unit (X-Cite Xylis, Opto Science), and a motorized stage. The sHMRG-based assay was performed using a solution containing dsSiR[1] (10 μM) as the internal standard, and the focus was adjusted using its fluorescence. Images were acquired in tile scan mode with perfect focus. The excitation and emission filters used were FITC (mirror = 510 nm, Ex. = 460-500 nm, Em. = 510-560 nm) and mCherry (mirror = 600 nm, Ex. = 550-590 nm, Em. = 608-683 nm), respectively.

**Image processing**

Images were processed using the GA3 module of NIS Elements software (Nikon). First, all fluorescence images were background-corrected using a rolling ball correction (3 μm). Then, ROIs were chosen by bright spot detection using mCherry filter settings (diameter = 3 μm), and irregular fluorescent spots derived from fluorescent debris or air bubbles were omitted by dilating the ROI and removing the overlapping ROIs. The fluorescence signals were calculated as the mean of the signals from the 9 pixels at the center of each ROI. Data were processed using Excel, or Kaleidagraph software to construct histograms and scatter plots.

**Plasma samples from mice**

Ethical approval for the study using animals was obtained from the Animal Care and Use Committee of The University of Tokyo (P4-21, P31-9). Six-week-old male C57BL/6JJcl mice were purchased from CLEA Japan (Tokyo, Japan) and acclimatized for five days. The mice were exposed to thioacetamide (TAA, T0817, Tokyo Chemical Industry Co., Japan, 350 mg/L) or 4,4'-methylene dianiline (MDA, M0220, Tokyo Chemical Industry Co., Japan, 750 mg/L) dissolved in drinking water to induce liver damage, whereas control mice received tap water. After four days of treatment, mice were euthanized and blood was collected from the inferior vena cava into 1.5 mL tubes containing 1.5 μL heparin (Yoshindo Inc, Japan). The collected blood sample was centrifuged (1,700 g, 4°C for 15 min) for plasma separation. Characterization of the blood samples and the detailed disease states of mice are shown in the reference[2].

### Supplementary methods for synthesis and characterization of compounds

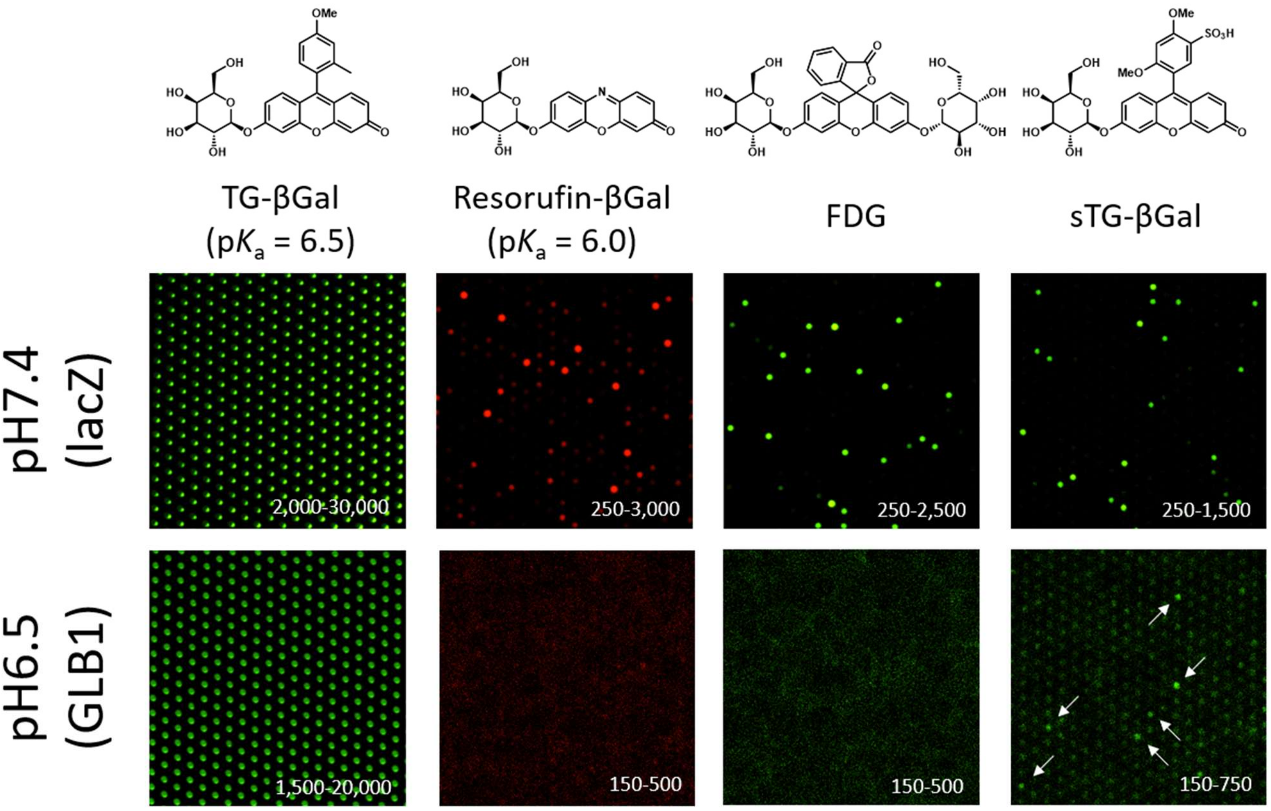


**Figure S1.** Suitability of β-galactosidase probes for single-molecule enzyme acvitiy assay. Probes (30 μM) mixed with β-galactosidase (β-galactosidase from *E. coli* (lacZ) for neutral and β-galactosidase from *Homo sapiens* (GLB1) for acidic) (1 ng/mL) in phosphate buffer (pH 7.4 or 6.5) containing Triton X-100 (30 μM) were loaded into microdevice and incubated at 25°C for 18 h.

**Supplementary methods for preparation of compounds**

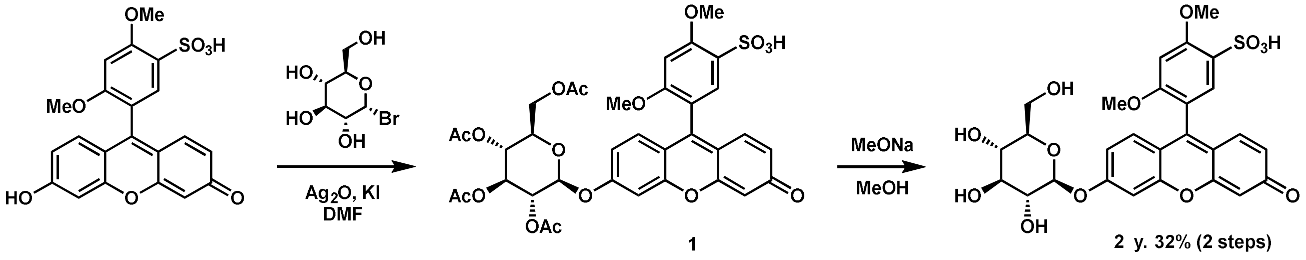


**Scheme S1.** Synthesis of sTG-based glycosidase probes with direct carbohydarate modification. The scheme for sTG-βGlc is shown, and sTG-βGal and sTG-βGlcNAc were prepared using the same scheme.

### Preparation of sTG β-Glc

sTG was synthesised according to the literature[3]. A solution of sTG (10 mg, 23.4 mmol, 1.0 eq.) in 300 μL of dry DMF was added to a flask charged with 2,3,4,6-tetra-*O*-acetyl-α-D-glucopyranosyl bromide (96 mg, 10 eq.), $Ag_2O$ (84 mg, 15 eq.), KI (32 mg, 8 eq.) and a small potion of $Na_2SO_4$. The reaction mixture was stirred at room temperature for 16 h, filtered and evaporated. The residue was dissolved in MeOH (300 μL) and 1 M NaOH aq. was added (300 μL). The mixture was stirred at room temperature for 1 h, after which the organic solvent was removed *in vacuo*. The residue was purified by HPLC (a linear gradient formed from 100 mM TFAA buffer and $CH_3CN$ 99%, $H_2O$ 1%: isocratic at 90/10 for 5 min., linear gradient to 10/90 in 15 min., then isocratic for 5 min) to afford 4.7 mg of pale orange powder after lyophilisation (32% in 2 steps).

$^1$H NMR (400 MHz, $CH_3OD$) δ 3.48-3.58 (m, 4H), 3.67 (m, 1H), 3.80 (s, 3H), 3.88 (s, 1H), 4.03 (s, 3H), 4.17 (m, 1H), 4.70 (s, 1H), 4.76 (s, 1H), 4.88 (s, 1H), 5.77 (s, 1H, *J* = 1.7 Hz), 6.08 (s, 1H), 6.22 (d, 1H, *J* = 1.8 Hz), 6.41-6.44 (m, 2H), 6.84 (s, 1H), 6.91 (s, 1H), 7.13 (d, 1H, *J* = 2.0 Hz), 7.71 (s, 1H).

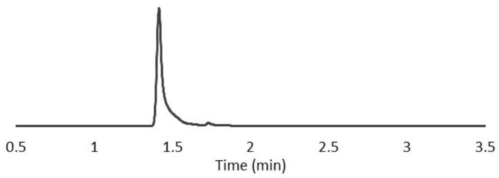


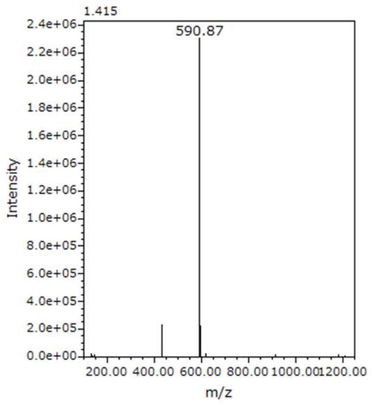


LC Chromatogram was monitored at 450 nm (A/B = 99/1 isocratic, 0.5 min, 99/1 to 0/100, 3.0 min; A = $H_2O$ + 0.1% TFA, B = AcCN-$H_2O$ (8:2) + 0.1% TFA).

LRMS (ESI$^+$): *m/z* = 591 (M+H)$^+$

### Preparation of sTG β-Gal

A solution of sTG (10 mg, 23.4 mmol, 1.0 eq.) in 300 μL of dry DMF was added to a flask charged with acetobromoα-D-galactose (99 mg, 10 eq.), $Ag_2O$ (84 mg, 15 eq.), KI (32 mg, 8 eq.) and a small potion of $Na_2SO_4$. The reaction mixture was stirred at room temperature for 16 h, filtered and evaporated. The residue was dissolved in MeOH (300 µL) and 1M NaOH aq. was added (300 µL). The mixture was stirred at room temperature for 1 h, after which the organic solvent was removed *in vacuo*. The residue was purified by HPLC (a linear gradient formed from 100 mM TFAA buffer and $CH_3CN$ 99%, $H_2O$ 1%: isocratic at 90/10 for 5 min., linear gradient to 10/90 in 15 min., then isocratic for 5 min.) to afford 2.1 mg of pale orange powder after lyophilisation (19% in 2 steps).

$^1$H NMR (400 MHz, $CH_3OD$) δ 3.51-3.60 (m, 4H), 3.68 (m, 1H), 3.80 (s, 3H), 3.89 (s, 1H), 4.03 (s, 3H), 4.18 (m, 1H), 4.67 (s, 1H), 4.74 (s, 1H), 4.85 (s, 1H), 5.76 (s, 1H, *J* = 1.8 Hz), 6.08 (s, 1H), 6.21 (d, 1H, *J* = 1.8 Hz), 6.40-6.44 (m, 2H), 6.84 (s, 1H), 6.92 (s, 1H), 7.13 (d, 1H, *J* = 2.1 Hz), 7.72 (s, 1H).

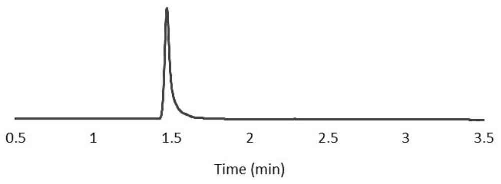


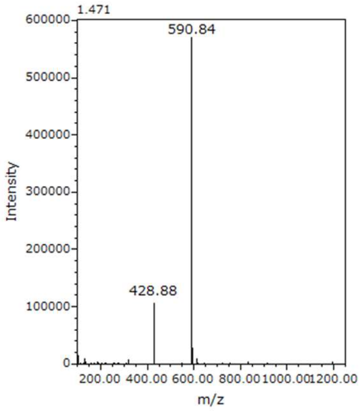


LC Chromatogram was monitored at 450 nm (A/B = 99/1 isocratic, 0.5 min, 99/1 to 0/100, 3.0 min; A = $H_2O$ + 0.1% TFA, B = AcCN-$H_2O$ (8:2) + 0.1% TFA).

LRMS (ESI$^+$): *m/z* = 591 (M+H)$^+$

**Preparation of sTG β-GlcNAc**

A solution of sTG (10 mg, 23.4 mmol, 1.0 eq.) in 300 µL of dry DMF was added to a flask charged with 2-acetamido-3,4,6-tri-*O*-acetyl-2-deoxy-α-D-glucopyranosyl chloride (89 mg, 10eq.), $Ag_2O$ (84 mg, 15 eq.), KI (32 mg, 8 eq.) and a small potion of $Na_2SO_4$. The reaction mixture was stirred at room temperature for 16 h, filtered and evaporated. The residue was dissolved in MeOH (300 µL) and 1M NaOH aq. was added (300 µL). The mixture was stirred at room temperature for 1 h, after which the organic solvent was removed *in vacuo*. The residue was purified by HPLC (a linear gradient formed from 100 mM TFAA buffer and $CH_3CN$ 99%, $H_2O$ 1%: isocratic at 90/10 for 5 min., linear gradient to 10/90 in 15 min., then isocratic for 5 min.) to afford 4.7 mg of pale orange powder after lyophilisation (32% in 2 steps).

$^1$H NMR (400 MHz, $CH_3OD$) δ 2.05 (s, 3H), 3.47-3.55 (m, 4H), 3.81 (s, 3H), 3.92 (s, 1H), 4.04 (s, 3H), 4.18 (m, 1H), 4.29-4.35 (m, 3H), 4.48 (s, 1H), 6.08 (s, 1H), 6.21 (d, 1H, *J* = 1.8 Hz), 6.39-6.45 (m, 3H), 6.83 (s, 1H), 6.88 (s, 1H), 7.13 (d, 1H, *J* = 2.1 Hz), 7.69 (s, 1H), 8.09 (s, 1H).

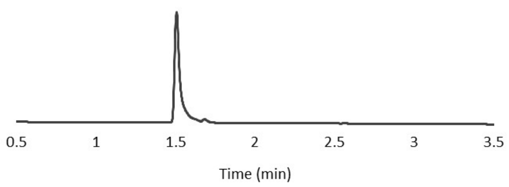


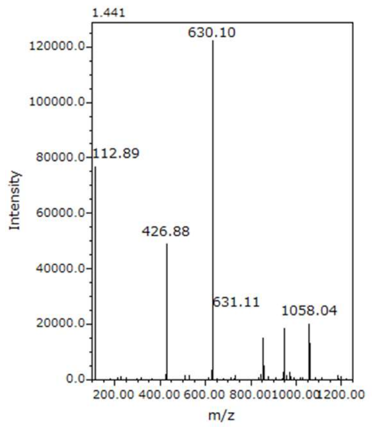


LC Chromatogram was monitored at 450 nm (A/B = 99/1 isocratic, 0.5 min, 99/1 to 0/100, 3.0 min; A = $H_2O$ + 0.1% TFA, B = AcCN-$H_2O$ (8:2) + 0.1% TFA).

LRMS (ESI$^-$): *m/z* = 630 (M+H)$^-$

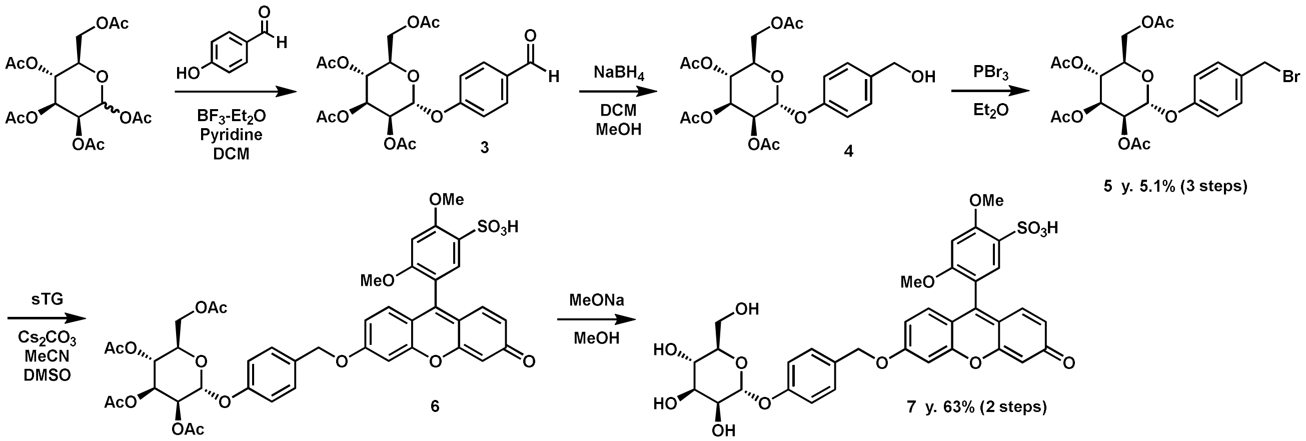


**Figure S2.** Synthesis of sTG-based glycosidase probes with direct carbohydarate modification (sTG-αMan).

**Preparation of compound 5.**

A mixture of *p*-benzaldehyde (1.10 g, 9.01 mmol), 1,2,3,4,6-Penta-*O*-acetyl-D-mannopyranose (3.50 g, 8.97 mmol), boron trifluoride-ethyl ether complex (8.0 mL), pyridine (2.0 mL) in 15 mL dried DCM was stirred at room temperature under an Ar atmosphere for 20 h. The reaction was quenched with sat. $NaHCO_3$ aq., the organic layer was washed with water, brine, dried over $Na_2SO_4$ and concentrated. The residue was purified by flash column chromatography (AcOEt : hexane = 0 : 100 to 50 : 50) to obtain the crude 1,2,3,4,6-penta-*O*-acetyl-D-mannopyranosylated derivative. The crude product was dissolved in a 20.0 mL solution of dried MeOH and DCM (MeOH : DCM = 1 : 1). To this solution, $NaBH_4$ (300 mg, 7.93 mmol) was added, and the reaction mixture was stirred at 0°C for 3 h. The reaction mixture was quenched with $NH_4Cl$ aq., extracted with DCM, dried over $Na_2SO_4$, and concentrated *in vacuo*. The residue obtained was dissolved in 20.0 mL $Et_2O$. To this solution, $PBr_3$ (390 μL) was added, and the reaction mixture was stirred at 0°C for 2 h. The reaction mixture was quenched with sat. $NaHCO_3$ aq., extracted with AcOEt, brine, dried over $Na_2SO_4$ and concentrated *in vacuo*. The residue was purified by flash column chromatography (AcOEt : hexane = 0 : 100 to 50 : 50) to afford compound **5** (236 mg, 0.457 mmol) as a white powder in 5.1 % yield. Identity of the compound was confirmed by comparing its spectroscopic data with those reported previously[4].

HRMS ($ESI^+$): *m/z* calcd. for $[M+Na]^+$, 577.17981; found, 577.18002.

**Preparation of sTG α-Man**

A mixture of sTG (17 mg, 0.04 mmol), compund **5** (100 mg, 0.19 mmol), $Cs_2CO_3$ (83 mg, 0.26 mmol) in a 5.0 mL solution of dried MeCN and DMSO (MeCN : DMSO = 3 : 2) was stirred at room temperature under an Ar atmosphere for 4 h. The reaction mixture was filtered and evaporated to remove MeCN. The residue was purified by HPLC to obtain orange powder, which was dissolved in 10.0 mL dried MeOH. To this solution, NaOMe (50 mg, 0.93 mmol) was added, and the reaction mixture was stirred at room temperature for 2 h then concentrated under reduced pressure. The residue was purified by HPLC to afford sTG-αMan (17.3 mg, 0.025 mmol) as an orange powder in 63% yield.

$^1H$ NMR (400 MHz, $CD_3OD$): δ 7.87 (d, *J* = 2.0 Hz, 1H), 7.78-7.83 (m, 2H), 7.69 (d, *J* = 2.0 Hz, 1H), 7.50 (d, *J* = 8.8 Hz, 2H), 7.34-7.370 (m, 2H), 7.25 (dd, *J* = 2.4 Hz, *J* = 9.6 Hz, 1H), 7.19 (d, *J* = 8.8 Hz, 2H), 7.0 (s, 1H), 5.52 (t, *J* = 2.0 Hz, *J* = 3.6 Hz, 1H), 5.40 (s, 2H) 4.14 (s, 3H), 4.02 (dd, *J* = 2.0 Hz, *J* = 3.2 Hz, 1H), 3.91 (dd, *J* = 3.2 Hz, *J* = 9.2 Hz, 1H), 3.84 (s, 1H), 3.69-3.80 (m, 3H), 3.56-3.61 (m, 1H).

$^{13}C$ NMR (101 MHz, DMSO-$d_6$): δ 174.6, 167.4, 160.3, 160.0, 159.3, 158.2, 157.5, 156.6, 133.2, 131.9, 131.1, 130.0, 128.8, 128.5, 122.8, 118.3, 117.5, 116.7, 116.4, 109.6, 102.7, 101.1, 98.7, 97.0, 75.0, 71.0, 70.6, 70.0, 66.6, 61.0, 56.2, 56.0.

HRMS ($ESI^-$): *m/z* calcd. for $[M-H]^-$, 695.14673; found, 695.14400.

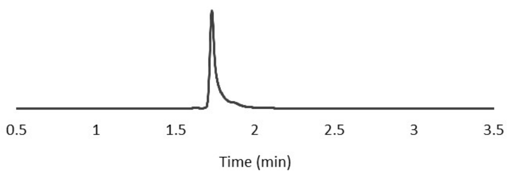


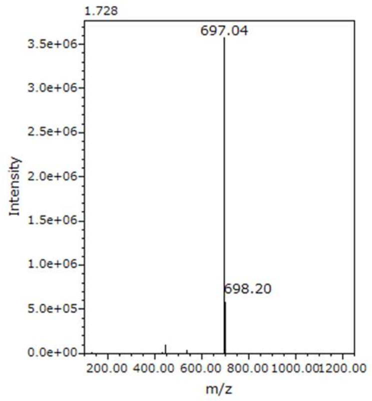


LC Chromatogram was monitored at 450 nm (A/B = 99/1 isocratic, 0.5 min, 99/1 to 0/100, 3.0 min; A = $H_2O$ + 0.1% TFA, B = AcCN-$H_2O$ (8:2) + 0.1% TFA).

LRMS ($ESI^+$): *m/z* = 697 $(M+H)^+$
